# Achieving the Endgame: Integrated NTD Case Searches

**DOI:** 10.1101/352336

**Authors:** Lucas Buyon, Randall Slaven, Paul M. Emerson, Jonathan King, Oscar Debrah, Agatha Aboe, Ernesto Ruiz-Tiben, Kelly Callahan

**Affiliations:** Harvard T.H. Chan School of Public Health, Department of Immunology and Infectious Diseases, Boston, MA, 02115, United States; The Carter Center, Atlanta, GA, 30307, United States.; International Trachoma Initiative, Decatur, GA, 30030, United States; World Health Organization, Geneva, Switzerland.; Eye Care Unit, Ghanaian Health Service, Accra, Ghana; Sightsavers International, Boston, MA, 02108, United States

## Abstract

Trachoma and Guinea Worm Disease (GWD) are neglected tropical diseases (NTDs) slated for elimination as a public health problem and eradication respectively by the World Health Organization. As these programs wind down, uncovering the last remaining cases becomes an urgent priority. In 2010, The Ghana Health Service, along with The Carter Center, Sightsavers, and other partners, conducted integrated case search for cases of both GWD and the last stage of trachoma disease, trachomatous trichiasis (TT), as well as providing treatment for trachoma to meet elimination and eradication targets. House to house case search for both diseases was conducted and two case management strategies were explored: a centralized referral to services method and a Point of Care (POC) delivery method. 835 suspected TT cases were discovered in the centralized method, of which 554 accepted surgery. 482 suspected TT cases were discovered in the POC method and all TT cases accepted surgery in the POC searches. The cost per TT case examined was lower in the POC searches compared to the centralized searches ($19.97 in the POC searches and $20.85 in the centralized searches). Both strategies resulted in high surgical uptake for TT surgery, with average uptakes of 72.4% and 83.9% for the centralized and POC searches respectively. We present here that house to house case search offering services at POC are feasible and a potential tool for elimination and eradication programs nearing their end.

**Author Summary:** Trachoma and Guinea Worm Disease (GWD) are neglected tropical diseases (NTDs) slated for elimination as a public health problem and eradication respectively by the World Health Organization. As these programs wind down, uncovering the last remaining cases becomes an urgent priority in order to confirm that eradiation/elimination targets have been reached. Active case searches are one method of finding these last vestiges of disease. Searches for that look for multiple diseases are referred to as integrated searches. We piloted here integrated case searches for GWD and Trachoma with two case management strategies, a referral approach to a central location, and point of care approach (POC). POC approaches can difficult to implement in low resource settings because they require extensive personnel, financial, and logistical, support. However, POC approaches remove one of the biggest barriers to treatment, time spent traveling to a health center, and thus can improve treatment uptake. We found here that integrated active cases searches with a POC case management approach can be implemented in a low resource setting; and improve acceptance and uptake of trachoma examination and trichiasis surgery respectively without costing much more than the referral case management approach.

## Introduction

The neglected tropical diseases (NTDs) are a diverse group of diseases that affect the poorest of the poor, primarily in rural Africa, Asia and Latin America.^1^ Eight NTDs are slated for elimination, and two, yaws and *Dracunculus medinensis*, or Guinea worm disease (GWD), are targeted for eradication by World Health Organization (WHO) resolutions WHA39.21^2^ and WHA 66.12.^3^ If eradication is successful, they will join smallpox and rinderpest as the only diseases eradicated from the world. Trachoma is scheduled for elimination as a public health problem by the year 2020 by the WHO resolution WHA 51.11.^4^

Guinea worm disease (GWD) is a parasitic disease. Individuals become infected by drinking from a stagnant water source that contains copepods which carry larva from the parasitic nematode roundworm *Dracunculiasis medinensis*.^5^ Once ingested, *D. medinensis* larva mature and mate inside the host’s body. Approximately one year later, a female worm will emerge through a blister, causing immense pain. Infected persons often seek relief from the burning sensation by immersing the wound in a water source, at which point the worm releases its larvae.^5^ There is no known medical treatment available to kill the parasitic nematode while it is inside the host, and on average GWD incapacitates its victims for about 8.5 weeks.^5 6^

Eradication efforts for GWD primarily use behavioral and environmental interventions to break the cycle of transmission to prevent contamination of drinking water sources and by improving access to clean drinking water^7^. The eradication of GWD requires that every case must be found and contained, making surveillance a priority for countries on the verge of elimination.

Trachoma is caused by a bacterium, *Chlamydia trachomatis*, and is spread via direct contact with ocular and nasal discharge from an infected person, including unwashed towels or clothes, as well as eye-seeking flies.^8^ Over time, repeated cycles of infection and resolution result in scar formation on the inner eyelid, which contract and physically distort the eyelid. Due to this scarring and distortion, the eyelashes turn inward and begin to scrape the globe of the eye, causing trauma to the cornea that can result in blindness.^8^ This stage is referred to as trachomatous trichiasis (TT). Control for trachoma utilizes the WHO endorsed SAFE strategy (surgery, antibiotics, facial cleanliness, and environmental improvement).^10^ Trachomatous trichiasis can be addressed via surgery, which consists of a simple eye-lid rotation to stop the lashes from scratching the globe of the eye. Surgery to reduce the prevalence of TT in a district is critical to achieving elimination targets set by WHO (less than one case of TT per one-thousand persons) which makes detecting and treating cases of TT a priority for trachoma programs as they reach elimination thresholds.

For countries in the end stage for both eradication of GWD and elimination of trachoma as a public health problem, or elimination of other NTDs, there exists an opportunity for integrating case searches for the diseases of interest. As GWD eradication and national efforts to eliminate trachoma draw closer to their elimination thresholds, detecting the remaining cases is paramount. Ghana is one such country that reached this stage of both eradication of GWD and trachoma elimination in early 2010. Prior to the start of the trachoma elimination campaign, the Ghana Health Services (GHS), the implementing agency of the Ministry of Health, had set a goal of treating all unaddressed cases of TT. This aggressive goal exceeded the elimination threshold (1 case/1000 persons) set by WHO, but would ensure that Ghana would meet the requirements of elimination and reduce any risk of disease progression later.^11 12^ In 2010 the GHS, assisted by The Carter Center and Sightsavers, collaborated to conduct house-to-house case searches to identify the remaining TT cases to help meet this threshold. Concurrently, Ghana was in the certification of elimination process for GWD, presenting an opportunity to develop integrated case searches for both diseases. Because the TT case searches were already being conducted, the GHS, The Carter Center, and SightSavers, realized that it would be feasible and likely cost efficient to pilot a program to integrate rumor investigation for GWD in these existing searches. This created the opportunity to determine if an active case search was a feasible strategy, and in this instance, to jointly uncover cases of TT and investigate rumors of GWD to help aid in the goal of elimination and eradication of these diseases.

House-to-house case searches for TT and GWD had not been previously integrated by the GHS. This type of search would ideally provide a clear picture of the disease burden in the region, but because such searches are labor intensive, they can be difficult to implement. Therefore, this presented an opportunity to assess the feasibility of integrated house-to-house case searches. Furthermore, since uptake of services to examine and treat TT are often low^12–14^, mostly due to logistical barriers and the economic burden resulting from these logistical barriers, the integrated case searches were coupled with either a centralized case management approach (where patients where referred to a central care facility) or point of care case management approach (POC, where patients were offered care either within or near their home). This would enable assessment of: 1) the capacity to integrate case finding for multiple diseases and 2) the impact of two case management strategies on TT examination and surgical uptake: centralized case management versus POC case management.

## Methods

The GHS identified 29 districts with a TT backlog requiring interventions such as surgery, epilation, or other follow-up. The case searches consisted of active house-to-house searching with teams walking from house-to-house to identify cases. Search teams were comprised of GHS staff, Carter Center staff, and village based health workers (VBHWs). Districts were mapped at the household level prior to the searches to identify villages, regional centers, and household locations, to ensure that no households were missed. The GHS was responsible for the primary ideation and strategy of the case searches, the overall management of the searches, and, in conjunction with The Carter Center, the training of the case search teams. The Carter Center was additionally responsible for the ideation of the integrated case searches, logistics, and supplies for the search teams. Sightsavers was involved with the training of the surgical teams and aids, as well assisting with reporting and financing the searches.

The TT case searches involved a house-to-house search with team members describing a case of TT to persons in the local language, and asking if they had seen such presentations. The GWD searches were conducted during the same household visits, with team members showing photos of non-emergent and emergent guinea worm, and also asking in the local language if they had seen such presentations. Teams would use chalk to mark households for TT or GWD to verify the case search was completed and the subsequent findings for further investigation.

The difference between the two searches was their geographic location and importantly case management approaches: the first, conducted in February 2010, utilized a passive community referral system and centralized surgical units, referred to henceforth as the centralized management approach, and the second, conducted in November 2010, offered point of care services with surgical services either in homes or within 0.5 km of walking distance (e.g. the center of a village), referred to henceforth as the Point of Care (POC) management approach.

For both the centralized and POC case management approaches, the number of GWD and TT cases investigated and confirmed were recorded, as well as the number of persons eligible for TT surgery, and the number accepting TT surgery. The cost of paying the healthcare workers and surgeons, as well as logistical and surgical costs were calculated by the hours for each professional and the number of surgeries ultimately performed. The costs generated represent the integrated searches, not solely for the cost of TT screening.

### Centralized Case Management Approach

The centralized case management approach was conducted in four districts: Savelugu, Tolon-Kumbungu, Gushegu, and Karaga. Searches were conducted by Guinea worm VBHWs, who were trained to identify TT cases as a group at a central location the day before the district wide searches began. A total of 959 health workers staffed this search, and the vast majority of the health workers were male.

The workflow for the search was as follows: All individuals identified exhibiting signs or symptoms consistent GWD were transported to a case containment center for verification. Persons with suspected TT were referred to a central location for examination and, if eligible for surgery, given a referral for surgery at that same location. In the event that a surgical team was not nearby, referred patients were told where and when to report for surgery, at an assigned hospital, clinic, or school within 10 kilometers of their village.

### Point of Care Case Searches

The POC case management approach focused on delivery of TT examination and surgery at the point of care and was conducted in two districts: Central Gonja and West Gonja. Each team increased numbers of volunteers relative to the centralized searches. The workflow for POC case management was as follows: Persons screened to have signs or symptoms consistent with TT were placed on a list of possible cases, and subsequently diagnosed by a healthcare professional in or near their home. As with the centralized case management approach, individuals identified exhibiting signs or symptoms consistent GWD were transported to a case containment center for verification. After diagnosis, patients were treated either in their own home or within 0.5 kilometers away if they consented to treatment. The POC case management approach used a mobile surgery team familiar with the local conditions. The surgical teams could perform TT surgery within the house of the TT patient and provide follow up care in the same location. In total, 1,833 people were involved in the POC case management approach. The Red Cross volunteer group, mostly women, were a significant majority of these human resources.

### Analysis

Basic calculations were conducted using Microsoft Excel. To calculate households reached per person days of effort, we summed the total number of households reached in each search. Then we multiplied the total number of volunteers in each of the searches by the number of days spent in district, and divided the total number of households reached by this number.

## Results

The summarized results of both case search strategies are presented in Figure 1. Figure 2 shows the direct comparison of the outcome metrics between two case management approaches.

**Fig 1.**
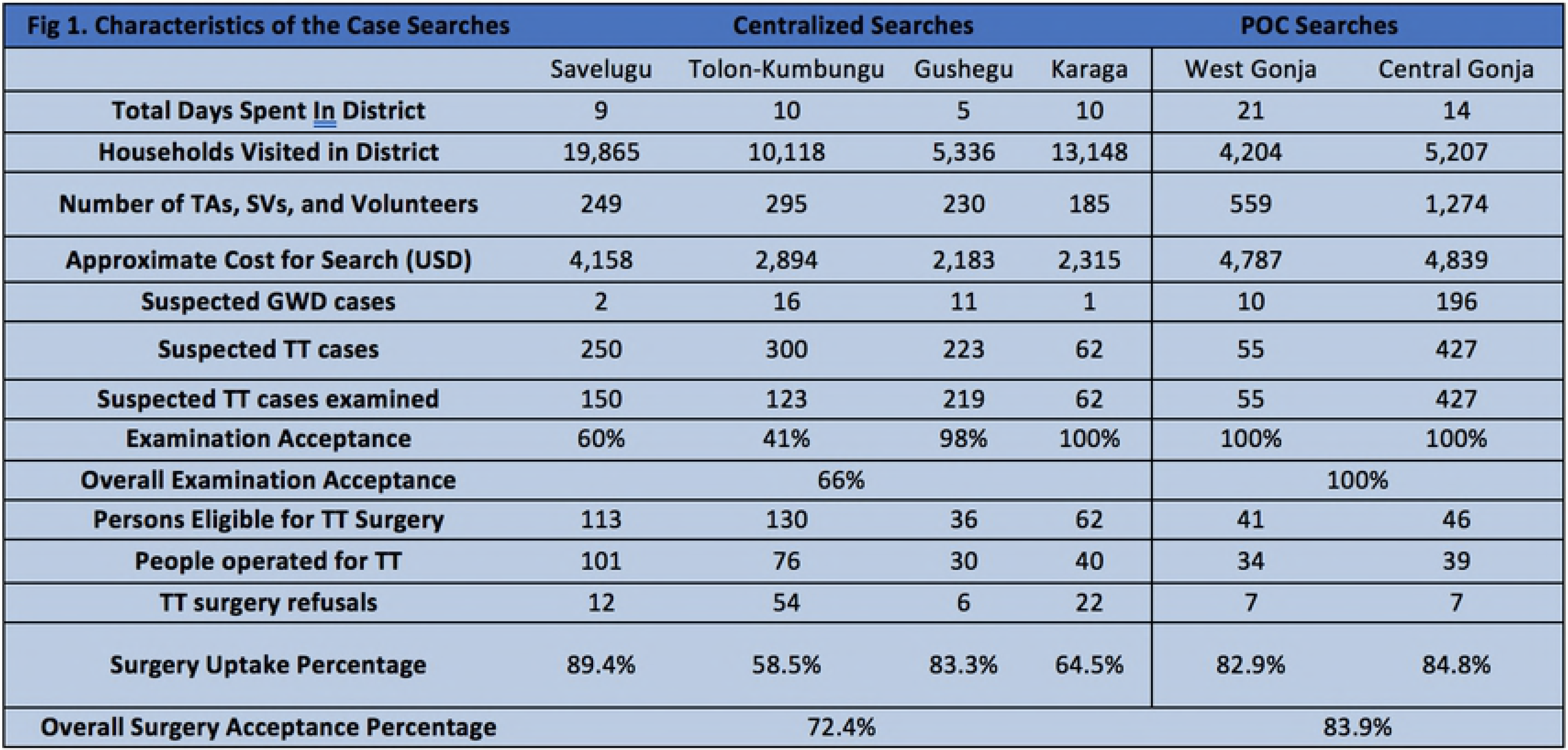
Table of characteristics and outcomes of the case searches

**Fig 2.**
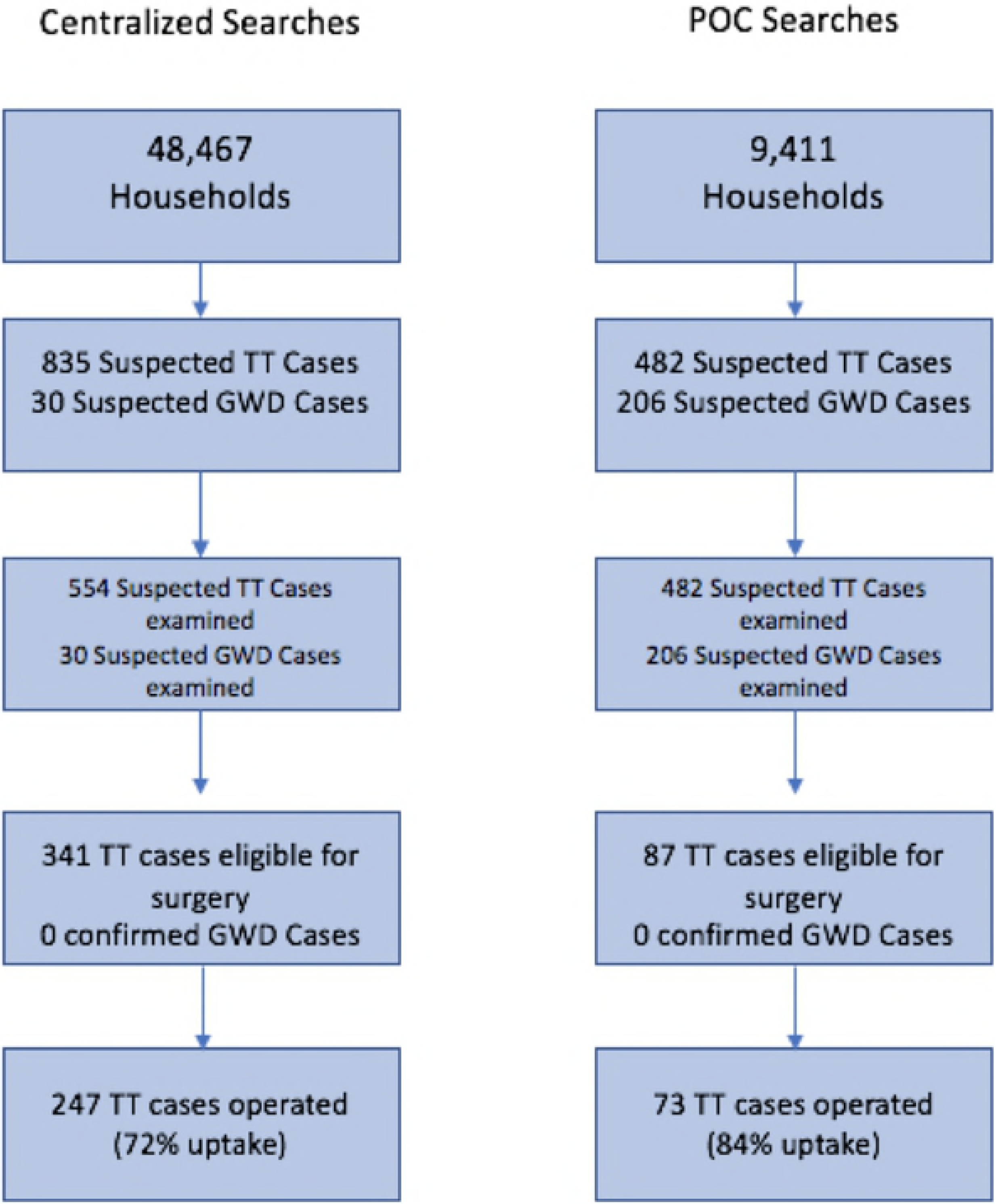
Flow chart contrasting results between methodologies

The centralized case management approach methodology reached 48,467 households using 959 volunteers and health workers over a period of 34 days. Accounting for the number of volunteers and days spent in district, this resulted in 1.49 households reached per person day of effort. The centralized case management approach also investigated a total of 30 individuals with signs or symptoms consistent with GWD. Additionally, 835 suspected TT cases were detected, for a rate of 17.2 TT cases suspected per 1,000 households reached. Of these, none were confirmed to be GWD and 554 TT suspected cases accepted examination (66.3% of cases found), of which 341 were eligible for surgery. The number of persons who accepted surgery was 247, resulting in an overall TT surgical uptake of 72.4% (247/341). The overall surgical refusal proportion was 27.6%. The TT surgical uptake percentage ranged from 58.5% - 89.4% across the four districts using the centralized case management searches. (Fig. 1)

The POC case management approach reached 9,411 households using 1,833 volunteers and health workers over a period of 35 days. Accounting for the number of volunteers and days spent in district, this resulted in 0.15 households reached per person day of effort. The POC searches investigated 206 individuals with signs or symptoms consistent with GWD. Furthermore, 482 TT cases were detected, for a rate of 51.2 cases of TT investigated per 1,000 households reached. Of these, none were confirmed to be GWD, and all 482 TT suspected cases accepted examination of which 87 were eligible for surgery. The number of persons who accepted surgery was 73, resulting in a surgical uptake of 83.9% (73/87). (Fig. 1)

The total cost of the centralized searches was $11,550 (excluding surgical interventions) or $0.24 per household visited and $20.85 per TT case examined. The total cost for the POC searches was $9,626 (excluding surgical interventions) or $1.02 per household visited and $19.97 per TT case examined.

## Discussion

Both case management approaches resulted in successful investigation of suspected TT and GWD cases, and achieved high TT surgical uptake, demonstrating that it is possible to a) conduct integrated case searches for multiple diseases, and b) deliver TT surgery at the point of care and achieve high TT examination and TT surgery uptake. In the both methods, the surgical uptake was high, 72.4% overall in the centralized case management approach and 83.9% overall in the POC case management approach. Notably, while TT examination acceptance was high in the centralized case management approach (41% to 100%, with an average of 66.3%), the acceptance was 100% in the POC searches. This likely resulted from POC approach removing the time investment and logistical barriers to undergo examination as patients received examination and care at a location convenient to them immediately.^12–14^ Additionally, an increase in the number of volunteers and the inclusion of women on every search teams likely improved TT examination uptake as well. Several previous studies have indicated that the biggest barriers to TT examination and surgical uptake are time lost to travel, cost of travel, and time spent on travel and recovery.^11–13^ For many of the world’s poor who live in rural areas, traveling to a central location for examination and surgery is a large investment—time spent traveling, waiting, and recovering equates to time not spent working, which creates an economic strain on the patient. The ability to deliver examination and surgery at the point of care removed these barriers by conducting examinations within or near patients’ homes, and doctors performing the surgery in a patient’s home. The results illustrate that offering TT examination and surgery at the point of care is not only feasible, but also potentially improves programmatic outcomes in regards to TT examination and surgical uptake.

In terms of cost, the centralized case management approach cost less on a per household basis ($0.24 per household visited) than the cost of the POC case management approach ($1.02 per household visited). The centralized case management approach was also more effective at reaching households compared to the POC case management approach (1.49 households reached per person day of effort compared to 0.15 households reached per person day of effort).

However, the rate of TT cases investigated per 1000 households visited was higher in the POC case management approach (50 cases of TT investigated per 1000 households reached) when compared to the centralized case management approach (17 cases of TT investigated per 1000 households reached). Furthermore, in terms of cost per TT case examined, the POC case management approach ($19.97 per case examined) cost slightly less than the centralized case management approach ($20.85 per case examined), since 100% of the TT cases investigated also accepted examination; likely due to the reasons stated in the previous paragraph.

This study had some limitations. Firstly, because this report and evaluation is observational in nature, and was not a formal trial, we cannot directly compare which approach is the most effective. Second, cultural and religious differences between the search regions could have also have contributed to differential examination acceptance rate. We also lacked data on the underlying levels of TT in these districts, which prevented us from gaging the direct impact of the searches on TT prevalence in these areas. Finally, we lacked previous data on TT surgical uptake rates in Ghana, and could not make a formal comparison as to how the current uptake compared to previous levels of surgical uptake. However, our aim here is not to evaluate the impact of the case search approach on TT prevalence, but rather to report and illustrate two ways of implementing case search approaches for TT, note the low cost and high TT examination acceptance and surgical uptake, and highlight the potential for integration with other NTD control and elimination programs.

The house-to-house case management approach offering examination and surgery at POC can be a workable and potent tool to meet eradication and elimination thresholds and help determine and relieve the burden of other NTDs. Many criticisms of point of care strategies stem from their relatively high cost when compared to centralized approaches. However, the fact that both methodologies presented here cost less than $12,000, reached tens of thousands of households, and achieved high TT surgical uptake, illustrates that integrated case searches using both methodologies are workable in the field. A POC case management approach might have been expected to cost more than a centralized case management approach due to the intensive logistical requirements. However, we demonstrate here that offering POC examination and surgery is feasible in a low income setting, achieves better services utilization, and costs slightly less relative to the centralized approach. The integrated house-to-house search and POC service approach could also be used to identify and treat unreported patients in need of care, such as the elderly, infirmed, and especially those afflicted with lymphatic filariasis and leprosy. The mixture of improved programmatic outcomes at similar or lower cost relative to a centralized approach warrants further exploration and usage of active case searches and POC approaches in low income settings for NTDs and other public health problems.

## Acknowledgements

We would like to thank Adam Weiss with his help and advice with manuscript.

## References

1. Hopkins, DR, Richards Jr FO, Ruiz-Tiben E, Emerson E, Withers PC. Dracunculiasis, Onchocerciasis, Schistosomiasis, and Trachoma. Annals of the New York Academy of Sciences. 2008; 1136(1): 45–52.

2. Hopkins, DR, Neglected Tropical Diseases slated for Elimination and Eradication. The Causes and Impacts of Neglected Tropical and Zoonotic Diseases: Opportunities for Integrated Intervention Strategies: Workshop Summary. 2001. National Academies Press.

3. World Health Organization (2013). WHA66.12 Neglected Tropical Diseases. Availible at: https://www.who.int/neglecteddiseases/mediacentre/WHA66.12Eng.pdf?ua=1. Accessed February 17, 2017.

4. World Health Organization (1998). "EB101.R5 Global elimination of blinding trachoma. Availible at: http://www.who.int/blindness/causes/EB101.R5/en/. Acessed February 17, 2017.

5. Muller, R. “Guinea worm disease: epidemiology, control, and treatment.” Bulletin of the World Health Organization. 1979; 57.5: 683.

6. Watts SJ, Brieger WR, Yacoob M. Guinea worm: an in-depth study of what happens to mothers, families and communities." Social Science & Medicine. 1989:29.9: 1043–1049.

7. Callahan K, Bolton B, Hopkins DR, Ruiz-Tiben E, Withers PC, Meagley K. “Contributions of the Guinea worm disease eradication campaign toward achievement of the Millennium Development Goals.” PLoS neglected tropical diseases 2013;7.5: e2160.

8. Mabey D C, Solomon AW, Foster A, Trachoma. Lancet. 2003; 362(9379): 223–229.

9. Thylefors, B, Dawson CR, Jones BR, West SK, Taylor HR. “A simple system for the assessment of trachoma and its complications.” Bulletin of the World Health Organaization. 1987; 65(4): 477–483.

10. World Health Organization (2017.) Trachoma: Status of endemicity for blinding trachoma, 2016. Availible at: https://rho.emro.who.int/rhodata/node.main.A1645?lang=en. Accessed 8/6/2017.

11. World Health Organization (1998). EB101.R5 Global elimination of blinding trachoma. Availible at: http://www.who.int/blindness/causes/EB101.R5/en/. Accessed February 17 2017.

12. Rajak SN, Habtamu E, Weiss HA, Bedri A, Zerihun M, Gebre T, et al. Why do people not attend for treatment for trachomatous trichiasis in Ethiopia? A study of barriers to surgery. PLoS Neglected Tropical Diseases. 2012;6(8): e1766.

13. Bowman, RJC, Faal H, Jatta B, Myatt M, Foster A, Johnson GJ, Bailey RL. Longitudinal study of trachomatous trichiasis in The Gambia: barriers to acceptance of surgery. Investigative Ophthalmology & Visual Science. 2002;43(4): 936–940.

14. Bowman, RJC, Sey Soma O, Alexander N, Milligan P, Rowley J, Faal H, Foster A, Bailey RL, Johnson GJ. Should Trichiasis Surgery be Offered in the Village? A Community Randomized Trial of Village vs. Health Centre-based Surgery. Tropical Medicine & International Health. 2000;5(8):528–533.

